# A chronic signaling TGFb zebrafish reporter identifies immune response in melanoma

**DOI:** 10.1101/2022.09.29.510035

**Authors:** Haley R. Noonan, Julia Barbano, Michael Xifaras, Chloé S. Baron, Song Yang, Katherine Koczirka, Alicia M. McConnell, Leonard I. Zon

**Affiliations:** Stem Cell Program and Division of Hematology/Oncology, Boston Children’s Hospital and Dana Farber Cancer Institute, Howard Hughes Medical Institute, Boston, MA 02115, USA.; Stem Cell and Regenerative Biology Department, Harvard University, Cambridge, MA 02138, USA.; Harvard Medical School, Boston, MA 02115, USA.; Biological and Biomedical Sciences Program, Harvard Medical School, Boston, Massachusetts 02115, USA.; Immunology Program, Harvard Medical School, Boston, Massachusetts 02115, USA.

## Abstract

Developmental signaling pathways associated with growth factors such as TGFb are commonly dysregulated in melanoma. Here we identified a human TGFb enhancer specifically activated in melanoma cells treated with TGFB1 ligand. We generated stable transgenic zebrafish with this TGFb Induced Enhancer driving green fluorescent protein (*TIE:EGFP*). *TIE:EGFP* was not expressed in normal melanocytes or early melanomas but was expressed in spatially distinct regions of advanced melanomas. Single cell RNA-sequencing revealed that *TIE:EGFP^+^* melanoma cells down-regulated interferon response, while up-regulating a novel set of chronic TGFb target genes. ChIP-sequencing demonstrated that AP-1 factor binding is required for activation of chronic TGFb response. Overexpression of SATB2, a chromatin remodeler associated with tumor spreading, showed activation of TGFb signaling in melanoma precursor zones and early melanomas. Confocal imaging and flow cytometric analysis showed that macrophages are recruited to *TIE:EGFP^+^*regions and preferentially phagocytose *TIE:EGFP^+^* cells. This work identifies a TGFb induced immune response and demonstrates the need for the development of chronic TGFb biomarkers to predict patient response to TGFb inhibitors.

## Introduction

Melanoma, arising from pigment producing melanocytes is the deadliest form of skin cancer, with an estimated 99,780 new cases and 7,650 deaths in the United States in 2022 alone^1^. The most common mutation in melanoma is BRAF^V600E^ which accounts for approximately 50% of melanoma cases and results in activation of the MAPK pathway promoting cell growth and survival^2, 3^. In addition, developmental signaling pathways are commonly dysregulated. Melanoma cells have increased expression and secretion of TGFb ligands compared to normal melanocytes, and TGFb ligand expression correlates with melanoma progression^4, 5,14,15, 6–13^. TGFb ligand binding to receptors on the cell surface results in phosphorylation and activation of SMAD2 and SMAD3 transcription factors. SMAD2 and 3 translocate to the nucleus with SMAD4 to modulate gene expression^16^. In normal melanocytes and early melanoma, TGFb acts as a tumor suppressor. However in advanced melanoma TGFb is pro-tumorigenic; inducing growth, invasion, and metastasis^10^. As current targeted MAPK and immune checkpoint inhibitors often result in resistance, there is a need to study additional pathways perturbed in melanoma, such as TGFb.

Most cells in the tumor microenvironment can respond to and initiate TGFb signaling, although this often occurs in a heterogenous manner^17^. Generally TGFb has an immunosuppressive effect, resulting in inactivation of cytotoxic CD8^+^ T-cells, expansion of immune suppressive regulatory T-cells, inhibition of dendritic cell antigen presentation, and conversion of macrophages to an anti-inflammatory and pro-angiogenic M2-like state^16–24^. TGFb can act as a chemoattractant for macrophages, recruiting monocytes to areas of inflammation. This results in differentiation into macrophages that attach to the ECM or promote blood vessel leakiness allowing for tumor cell extravasion^25–29^. In colorectal and urothelial cancers, TGFb in the tumor microenvironment was found to mediate immune evasion such that TGFb inhibition rendered these tumors susceptible to anti-PD-1-PD-L1 immune checkpoint inhibitors^30, 31^. Due to its immunosuppressive effect, several TGFb inhibitors are in clinical trials in combination with immune checkpoint inhibitors^32–36^.

Here, we visualized TGFb response across zebrafish melanoma tumorigenesis. A human melanoma enhancer, induced upon TGFB1 signaling was identified and stable transgenic zebrafish with this enhancer driving EGFP were generated (*TIE:EGFP*). This TGFb inducible enhancer reporter is expressed in spatially distinct regions in advanced zebrafish melanomas and is characterized by up-regulation of a series of novel chronic TGFb target genes involved in the extracellular matrix. Single cell RNA-seq and confocal microscopy revealed that *TIE:EGFP^+^* melanoma cells are preferentially phagocytosed by macrophages and downregulate interferon response. Overexpression of the chromatin remodeler SATB2, which is associated with tumor spreading, shows early activation of TGFb signaling in these melanomas, suggesting that specific melanoma genotypes may benefit from TGFb inhibition. Overall, this work demonstrates the need for biomarker development to predict response to TGFb inhibitors in advanced or aggressive melanoma subtypes.

## Results

### A TGFb Enhancer Reporter is inducible and specific in zebrafish

To visualize dynamic TGFb response across melanoma development, we designed a TGFb Inducible Enhancer (TIE) reporter using human melanoma cell (A375) ChIP-seq data following a 2-hour TGFB1 treatment. RNA-sequencing indicated that following a 2-hour treatment of A375 cells with 10ng/mL human recombinant TGFB1, 223 genes were significantly up-regulated (q<0.05) and 94 genes were down-regulated (Figure S1A left). This includes typical TGFb target genes such as SMAD7, JUNB, and PMEPA1. Hallmark gene set enrichment analysis (GSEA) of genes ranked by log2 fold-change (log2fc) confirmed that TGFb was the top up-regulated pathway following 2-hour treatment (q=0) (Figure S1A right). This indicates that a 2-hour treatment with human recombinant TGFB1 ligand is sufficient to activate the TGFb pathway.

To identify a TGFB1 inducible enhancer region we selected a region of chromatin that was exclusively open upon stimulation, based on H3K27ac ChIP-seq, with unique SMAD2/3 binding following treatment (Figure 1A). The enhancer region under the SMAD2/3 peak was cloned upstream of a beta-globin minimal promoter and EGFP in a Tol2 vector backbone. *TIE:EGFP* was tested for inducibility in the presence of ubiquitously expressed constitutively active SMAD2 and SMAD3 (*ubi:caSMAD2/3*) by electroporation of zebrafish skin, with ubiquitous BFP (*ubi:BFP*) as a control for electroporation efficiency. TIE reporter activity was significantly increased in the presence of *ubi:caSMAD2* and *ubi:caSMAD3,* confirming that the reporter is inducible (Extended Data Figure 1A).

**Figure 1.**
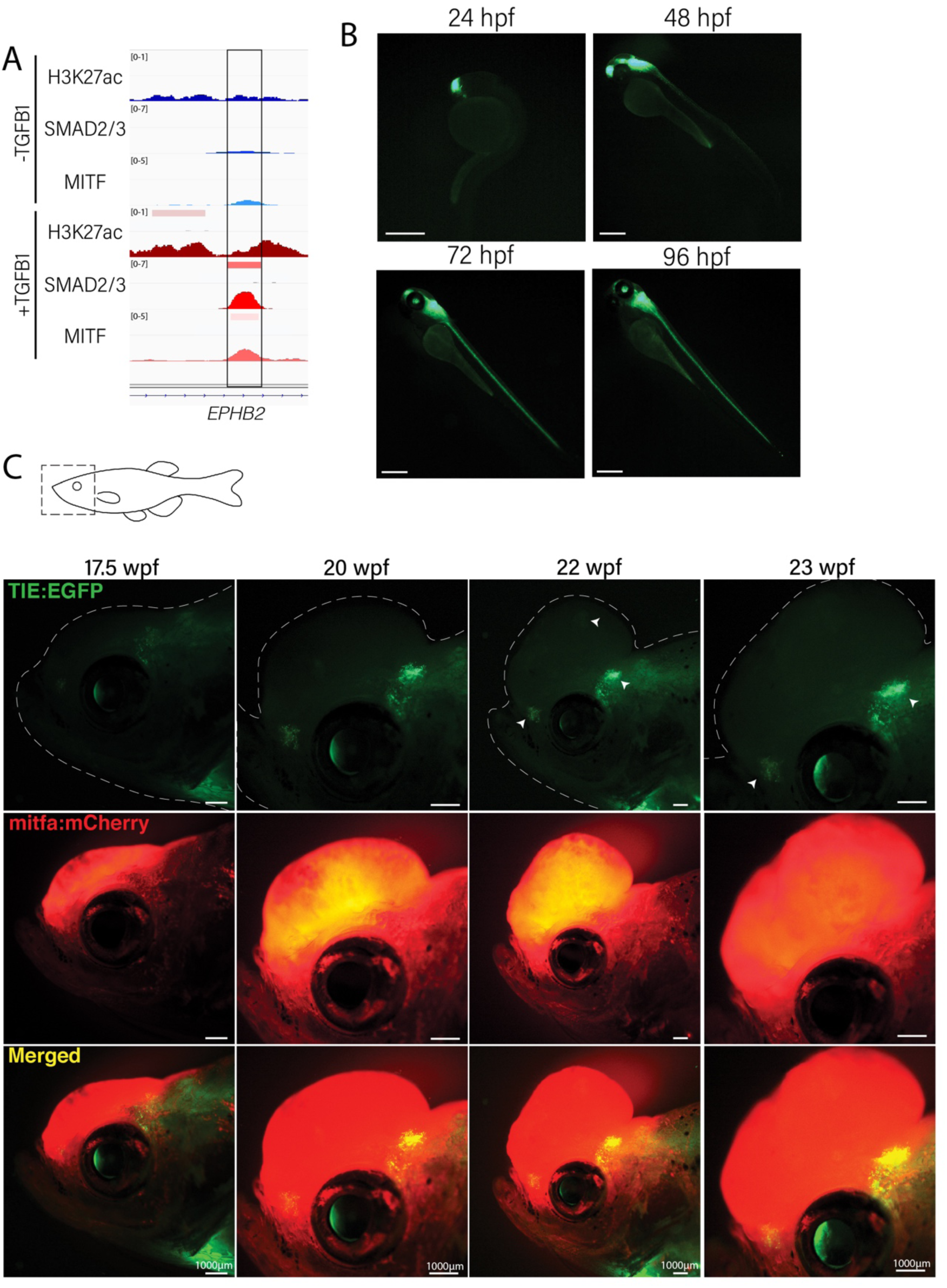
Novel *TIE:EGFP* zebrafish enhancer reporter is expressed in advanced melanomas. (A) TGFb induced enhancer (TIE) reporter H3K27ac, SMAD2/3, and MITF ChIP-seq peaks in A375s+/-TGFB1. There is unique H3K27ac and SMAD2/3 binding upon stimulus. (B) *TIE:EGFP* expression throughout zebrafish development. Scale bars represent 500µm. (C) *TIE:EGFP* expression across melanomagenesis indicated by arrowheads. Representative images shown.

**Extended Data 1.**
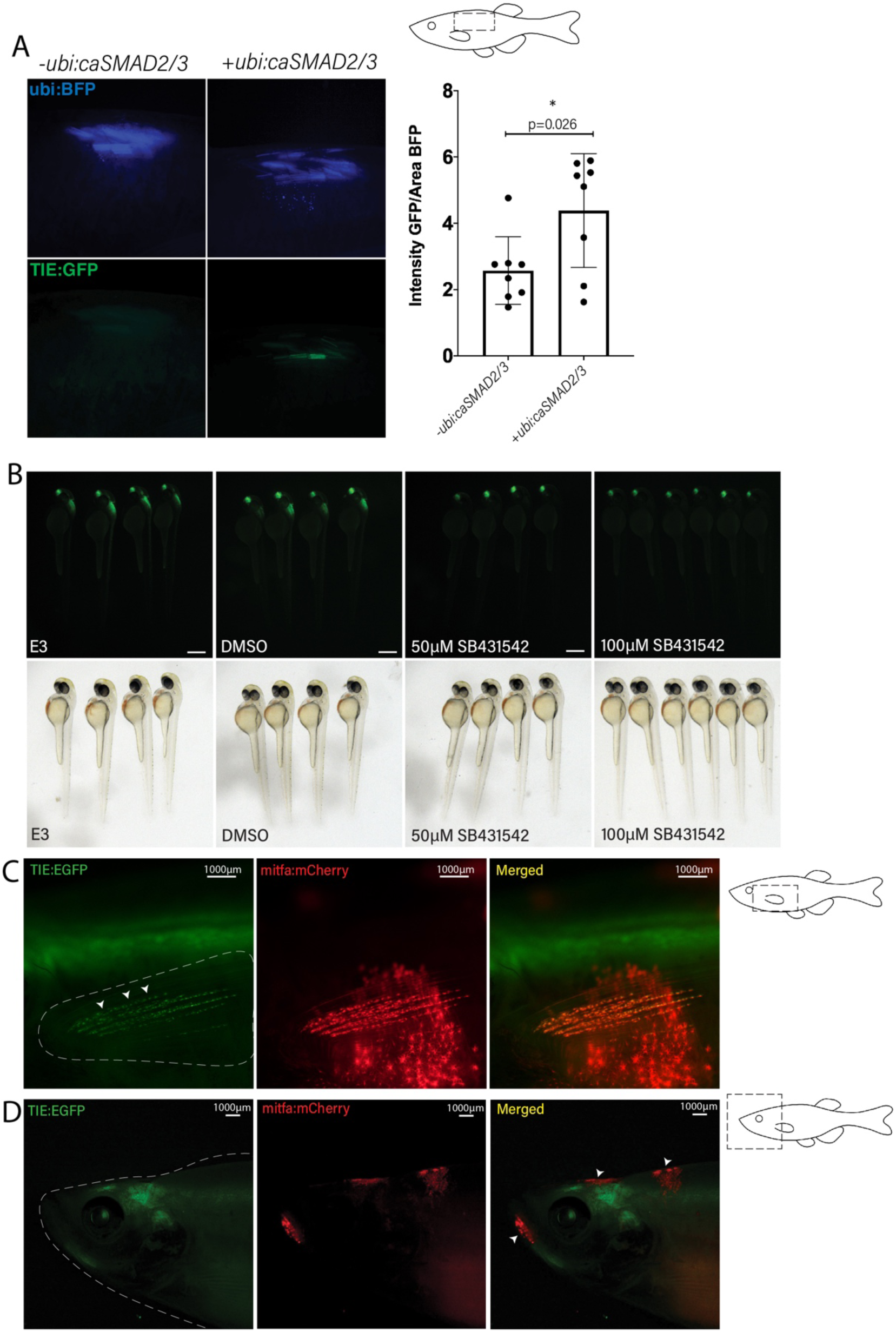
*TIE:EGFP* zebrafish enhancer reporter is inducible and specific. (A) *TIE:EGFP* reporter is induced upon electroporation with *ubi:caSMAD2* and *ubi:caSMAD3* in adult zebrafish flank skin. *Ubi:BFP* was used as a control for varying electroporation efficiency. *TIE:EGFP* Intensity/BFP area was quantified (right). Two-tailed unpaired Welch’s t-test was used to calculate significance. Representative images shown. n=8 fish per condition. (B) *TIE:EGFP* embryos treated at 1 day post fertilization for 24 hours with E3 (zebrafish water), DMSO (vehicle control), 50µM or 100µM TGFb inhibitor SB431542. *TIE:EGFP* channel (top) and white light (bottom). Representative images shown. Scale bars represent 500µm. (C) Representative image of *mitfa:mCherry*^+^ zebrafish fin melanocytes expressing *TIE:EGFP*. (D) Early melanomas indicated by arrowheads, do not express *TIE:EGFP*. Representative images.

The *TIE:EGFP* reporter was microinjected into *Tg(mitf:BRAF^V600E^); p53^−/−^; mitfa^−/−^* zebrafish in order to generate melanomas*. TIE:EGFP;Tg(mitf:BRAF^V600E^);p53^−/−^;mitfa^−/−^* stable embryos have EGFP expression in the anterior brain at 24 hours post fertilization (hpf) extending along the brain and spinal cord beginning at 48 hpf (Figure 1B). This expression pattern is consistent with that previously found in a Smad3 zebrafish reporter, although in our hands this Smad3 reporter was not active in adults^37^. *TIE:EGFP* embryos were treated at 24 hpf with 50 or 100 uM of SB431542, a TGFb type I receptor kinase inhibitor, and imaged at 48 hpf^27^. Treatment with TGFb inhibitor for 24 hours drastically reduced *TIE:EGFP* signal (Extended Data Figure 1B). This indicates that the *TIE:EGFP* reporter is specific to TGFb signaling.

### *TIE:EGFP* reporter is expressed in advanced zebrafish melanomas

To visualize TGFb response across melanoma development, single cell *Tg(mitf:BRAF^V600E^); p53^−/−^; mitfa^−/−^* embryos stably expressing *TIE:EGFP* were injected with an empty MiniCoopR vector (*MCR:MCS*) containing the *mitfa* minigene to generate melanomas. Tyrosinase gRNA and Cas9 protein was also injected to knock out melanocyte pigment as well as *mitfa:mCherry* to allow for melanocyte visualization. Normal melanocytes rarely had *TIE:EGFP* expression, with the exception of occasional fin melanocytes (Extended Data Figure 1C). *TIE:EGFP* was also never expressed in *mitfa:mCherry* high early melanomas (Extended Data Figure 1D). However, of the fish with melanoma formation (n= 56), 55% turned on *TIE:EGFP* in advanced melanomas defined by protrusion from the body plane (Figure 1C). *TIE:EGFP^+^* cells often occur in clusters throughout the tumor and these cells are *TIE:EGFP^+^* for many weeks at a time. This data indicates that most advanced zebrafish melanomas develop TGFb responsive zones.

### *TIE:EGFP* expressing melanoma cells downregulate interferon signaling

To understand the transcriptional differences between *TIE:EGFP^+^*and *TIE:EGFP^-^* melanoma cells, single *TIE:EGFP^+^*and *mitfa:mCherry^+^* cells from *TIE:EGFP* expressing melanomas were processed for single cell RNA-sequencing at 23 and 42 weeks post fertilization (wpf), respectively (Figure 1C and S1B). We used SORT-seq to sequence flow cytometry-sorted single cells and *in silico* linked transcriptomes to fluorescence intensities as recorded by FACS indexing^38^. A UMAP of the sequencing results from our combined replicates is shown (Figure 2A) (UMAP with no batch correction in Supplemental Figure 2A). Most cells are identified as melanoma cells expressing *mitfa* and/or *sox10*, but we identified a *marco*-expressing *TIE:EGFP^+^* macrophage cluster as well (Figure 2A).

**Figure 2.**
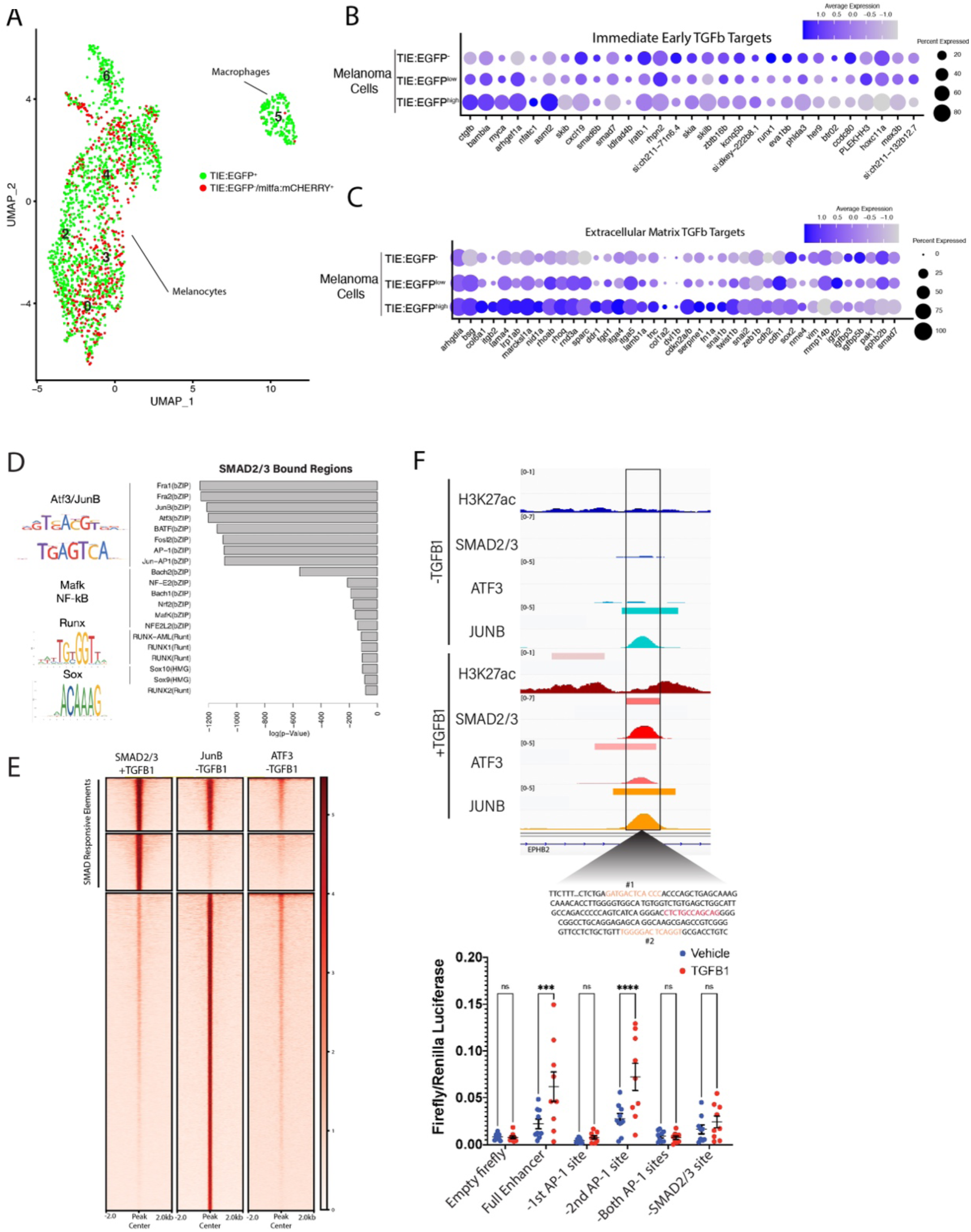
TGFb expressing *MCR:MCS* melanoma cells up-regulate chronic extracellular matrix TGFb target genes and AP-1 binding is required for TGFb responsiveness. (A) UMAP depicting 7 cell clusters identified by SORT-seq, combined 2 *MCR:MCS* replicates (left). Approximately 2,256 *TIE:EGFP^+^* cells and 752 *mitfa:mCherry^+^* cells were sorted for scRNA-seq. *TIE:EGFP^+^*cells were both mCherry^+^ and mCherry^-^. *TIE:EGFP^+^* and *TIE:EGFP^-^* cells are identified based on flow cytometry. Melanoma cells were identified as being *mitfa* and *sox10* positive, while macrophages are *marco* positive. (B) Dotplot depicting TGFb immediate early target gene expression in *TIE:EGFP^high^, TIE:EGFP^low^*, and *TIE:EGFP^-^* melanoma cells. Melanoma cells can be segregated into *TIE:EGFP^high^* vs*. TIE:EGFP^low^* based on EGFP intensity during sorting. (C) Dotplot depicting extracellular matrix TGFb target gene expression in *TIE:EGFP^high^, TIE:EGFP^low^*, and *TIE:EGFP^-^* melanoma cells. (D) HOMER motif analysis of regulatory regions bound by SMAD2/3 upon stimulation in A375 cells. (E) Heatmap showing binding of JUNB and ATF3 pre-stimulus at ∼12,000 SMAD2/3 responsive elements in A375. (F) Top: IGV plot of H3K27ac, SMAD2/3, ATF3 and JUNB ChIP-seq +/- TGFB1 stimulus at the TGFb-induced enhancer. Inset depicts sequence under SMAD2/3 ChIP-seq peak and highlights AP-1 (orange) and SMAD2/3 (red) binding sites. Bottom: Firefly luciferase luminescence of full TIE reporter or reporter lacking AP-1 or SMAD2/3 sites. Normalized to Renilla transfection control. Experiment performed 3 times with 3 technical replicates each. A 2-way multiple comparison ANOVA was used to calculate significance.

The melanoma cell population was separated into *TIE:EGFP^-^, TIE:EGFP^low^,* and *TIE:EGFP^high^* based on FACS intensities. We found that immediate-early TGFb target genes, identified following an acute 2-hour TGFB1 treatment in our zebrafish melanoma cells, ZMEL1, were down-regulated in *TIE:EGFP^high^* cells (Figure 2B). However, we identified 29 genes that were up-regulated in our *TIE:EGFP^high^* cells. We termed these genes, related to the extracellular matrix, “chronic” TGFb targets (Figure 2C)^39, 40^. At the time of dissociation, these melanomas had *TIE:EGFP^+^* cells for several weeks, therefore they were likely not in an acute TGFb response phase (Figure 1C and S1B). Using GSEA analysis we found that the top down-regulated pathways in *TIE:EGFP^high^* cells were interferon alpha and gamma (Figure S2B). This suggests that TGFb has an immune suppressive effect in our melanomas.

### AP-1 factors are required for induction of the chronic TIE reporter

Although the vast majority of literature focuses on acute TGFb signaling, our *TIE:EGFP* reporter remains on in melanoma, reading out chronic TGFb signal. We next asked what transcription factors are necessary for induction of our novel chronic TGFb reporter and therefore required for chronic TGFb signaling. We performed HOMER motif analysis of ∼12,000 SMAD2/3 responsive regulatory elements. These regions were identified in A375s melanoma cells stimulated with TGFB1 and were bound by SMAD2/3 upon stimulus. We found that these SMAD2/3 bound regions are highly enriched for AP-1 motifs (Figure 2D). ChIP-seq in A375 cells in the presence or absence of TGFB1 treatment confirmed the binding of AP-1 transcription factors JUNB and ATF3 (Figure 2E). ATF3 and JUNB were bound at 19% and 48% of SMAD2/3 responsive elements, respectively, before TGFB1 treatment was administered. This data indicates that AP-1 factors, in particular JUNB, may be important for SMAD binding to chromatin.

We hypothesized that AP-1 factors occupy SMAD responsive elements prior to stimulation, stabilizing open chromatin to allow the TGFb response to occur quickly upon stimulation. To test this hypothesis, we deleted AP-1 motifs in our TIE reporter driving luciferase. We identified AP-1 motifs (highlighted in orange) in our TGFb-induced enhancer using MoLoTool and HOCOMOCO motifs (Figure 2F top)^41^. AP-1 motif #1 was the most significant (p=8.4e^-6^), and AP-1 motif #2 had a lesser p-value of 8.2e^-4^. We deleted either AP-1 site #1 alone, AP-1 site #2 alone, or both AP-1 sites, as well as the most significant SMAD2/3 motif (p=6.7e^-6^), shown in red (Figure S3). In A375 cells, loss of AP-1 site #1 rendered our enhancer region unresponsive to TGFb signaling, however loss of AP-1 site #2 did not alter inducibility (Figure 2F bottom). This result indicates that AP-1 site #1 is necessary for TGFb inducibility at this enhancer. Deletion of the SMAD site also destroyed TGFB1 responsiveness. This data suggests that AP-1 binding is required for responsiveness of our chronic TGFb inducible enhancer in melanoma.

### Macrophages preferentially phagocytose *TIE:EGFP* expressing cells

We identified *TIE:EGFP^+^* macrophages in *TIE:EGFP*;*Tg(mitf:BRAF^V^*^600^*^E^);p53^−/−^;mitfa^−/−^* melanomas that express *mitfa* and *sox10,* suggesting recent phagocytosis of melanoma cells (Figure 3A). There are two potential models for why macrophages are *TIE:EGFP^+^*: (1) macrophages express TGFb themselves, or (2) macrophages are engulfing *TIE:EGFP*^+^ cells. To test these hypotheses, we crossed our *TIE:EGFP;Tg(mitf:BRAF^V^*^600^*^E^); p53^−/−^; mitfa^−/−^* zebrafish to a *mpeg:mCherry* reporter line that labels macrophages with mCherry^42^. These embryos were injected with *MCR:BRAF^V^*^600^*^E^, 2x:U6 p53/Tyr gRNA mitfa:Cas9*, and *mitfa:BFP* to generate pigmentless, BFP^+^ melanomas. As the *TIE:EGFP* reporter is cytoplasmic, if macrophages were endogenously expressing *TIE:EGFP*, we would expect the entire macrophage to be yellow. However, macrophages consistently contained only puncta of *TIE:EGFP* signal and we observed macrophages actively phagocytosing *TIE:EGFP^+^* cells (Figure 3B and S4). This data shows that macrophages are *TIE:EGFP^+^* because they are engulfing *TIE:EGFP^+^* tumor cells.

**Figure 3.**
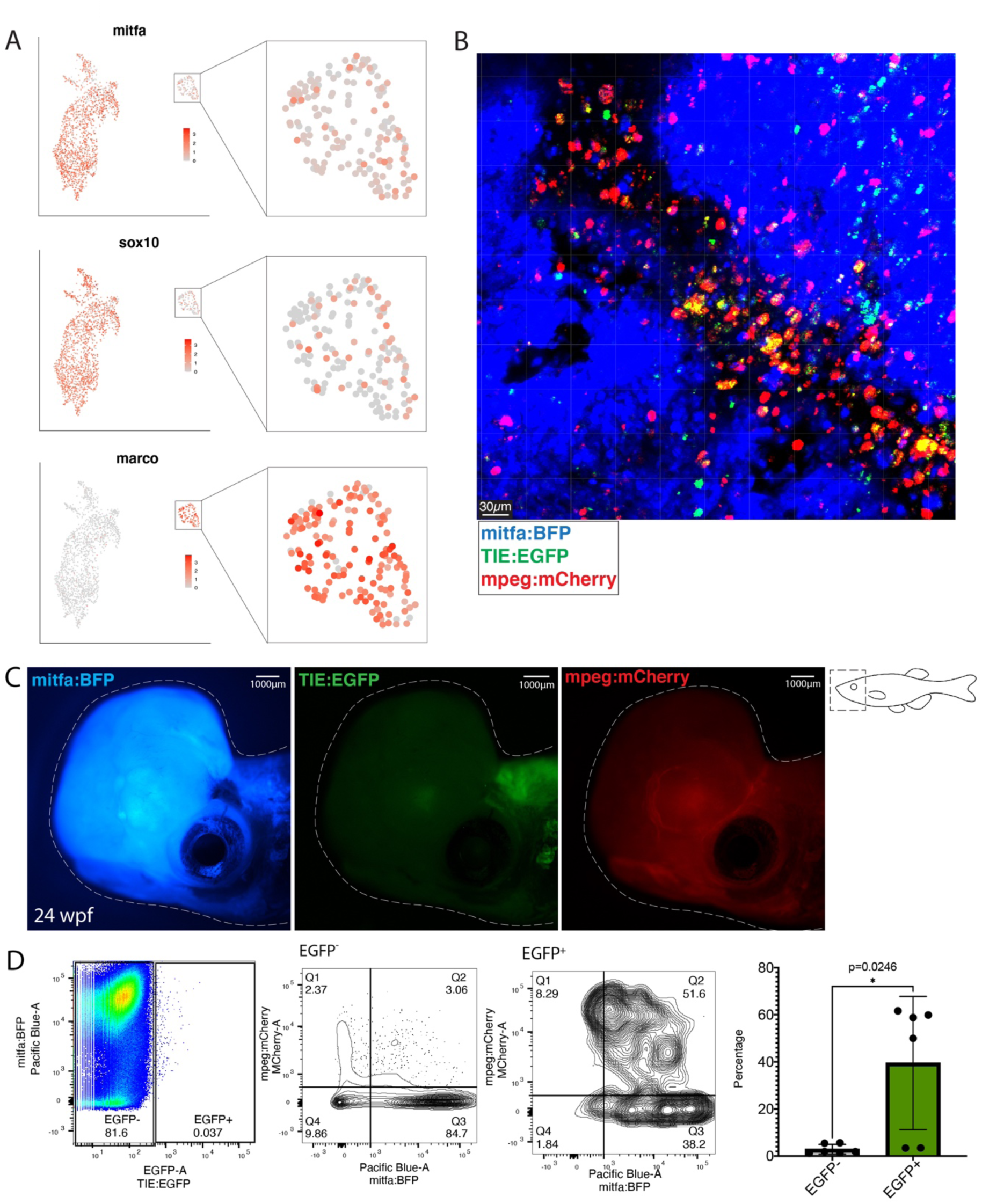
Macrophages preferentially phagocytose *TIE:EGFP^+^* cells. (A) UMAP depicting *mitfa, sox10,* and *marco* expression in clusters identified by SORT-seq, combined 2 *MCR:MCS* replicates. Inset shows expression of these genes in the macrophage cluster. (B) Representative image from a zebrafish melanoma acquired on an upright confocal, n=5 fish. Melanoma is blue, macrophages are red, and *TIE:EGFP^+^* cells are green. Yellow indicates a macrophage that has phagocytosed a TGFb positive non-melanoma cell. Cyan indicates a *TIE:EGFP^+^* melanoma cell. When phagocytosed by macrophages, *TIE:EGFP*^+^ melanoma cells appear white. (C) Representative *TIE:EGFP^+^*;*mpeg:mCherry^+^;mitfa:BRAF^V600E^;2x:U6 p53/Tyr gRNA mitfa:Cas9*;*mitfa:BFP*^+^ melanoma for flow analysis of macrophages. Scale bars indicate 1000µm. (D) Left: Viable cells were separated into *TIE:EGFP^-^* and *TIE:EGFP^+^*. Middle: FACs plots showing *TIE:EGFP^-^* and *TIE:EGFP^+^*cells relative to *mpeg:mCherry* and *mitfa:BFP*. Q1 and Q2 represent *TIE:EGFP*^-^ cells that were phagocytosed by macrophages. Q2 contains BFP^+^ melanoma cells that were phagocytosed, while Q1 contains non-melanoma cells that were phagocytosed. Right: The percentages in Q1 and Q2 were summed, to represent the total percentage of *TIE:EGFP*^-^ or *TIE:EGFP^+^* cells phagocytosed by macrophages. Two-tailed unpaired Welch’s t- test was used to calculate significance. n=3 fish with two technical replicates each.

To determine if macrophages are preferentially phagocytosing *TIE:EGFP*^+^ cells, three *TIE:EGFP^+^;mpeg:mCherry^+^;mitfa:BFP^+^*melanomas were excised, digested, and processed for flow cytometry analysis (Figure 3C). Viable cells were separated into *TIE:EGFP^-^* and *TIE:EGFP^+^* (Figure 3D left). In all three tumors, less than 1% of sorted cells were EGFP^+^ indicating this TGFb expressing population is rare. *TIE:EGFP*^-^ and *TIE:EGFP^+^* (Figure 3D middle) cells were plotted relative to *mpeg:mCherry* and *mitfa:BFP*. Q1 and Q2 represent *TIE:EGFP*^-^ or *TIE:EGFP^+^* cells that were phagocytosed by macrophages. Q2 contains BFP^+^ melanoma cells that were phagocytosed, while Q1 contains non-melanoma cells that were phagocytosed. The percentages in Q1 and Q2 were summed, to represent the total percentage of *TIE:EGFP*^-^ or *TIE:EGFP^+^* cells phagocytosed by macrophages and plotted in Figure 3D (right). Macrophages phagocytose a significantly higher percentage of *TIE:EGFP^+^* cells compared to *TIE:EGFP*^-^ cells, this is true for both *TIE:EGFP^+^* melanoma and non-melanoma cells. Altogether scRNA-seq, confocal imaging and flow cytometry analyses demonstrate that macrophages preferentially phagocytose *TIE:EGFP^+^*melanoma cells.

### SATB2 expression leads to early TGFb activation

The chromatin remodeler *SATB2* is amplified in 4-8% of melanoma patients and its expression correlates with patient outcome. *SATB2* expression leads to more aggressive, metastatic melanoma and induces a TGFb signature^43^. *SATB2* was overexpressed under the *mitfa* promoter in the MiniCoopR backbone (*MCR:SATB2*) in our *TIE:EGFP*;*Tg(mitf:BRAF^V600E^); p53^−/−^; mitfa^−/−^* zebrafish. Of the fish with SATB2 melanomas, 68% turned on *TIE:EGFP* (n=27) in tumors (Figure 4A). To understand the transcriptional differences between *TIE:EGFP^+^* and *TIE:EGFP^-^* melanoma cells, we sequenced the transcriptome of single cells from a *MCR:SATB2* expressing melanoma at 30 wpf using SORT-seq (Figure 4A). We identified most cells as melanoma cells with *mitfa* and/or *sox10* expression, and again a *marco-*expressing *TIE:EGFP^+^* population reproducing our previous finding of macrophages phagocytosing *TIE:EGFP^+^* cells (Figure 4B and S5A).

**Figure 4.**
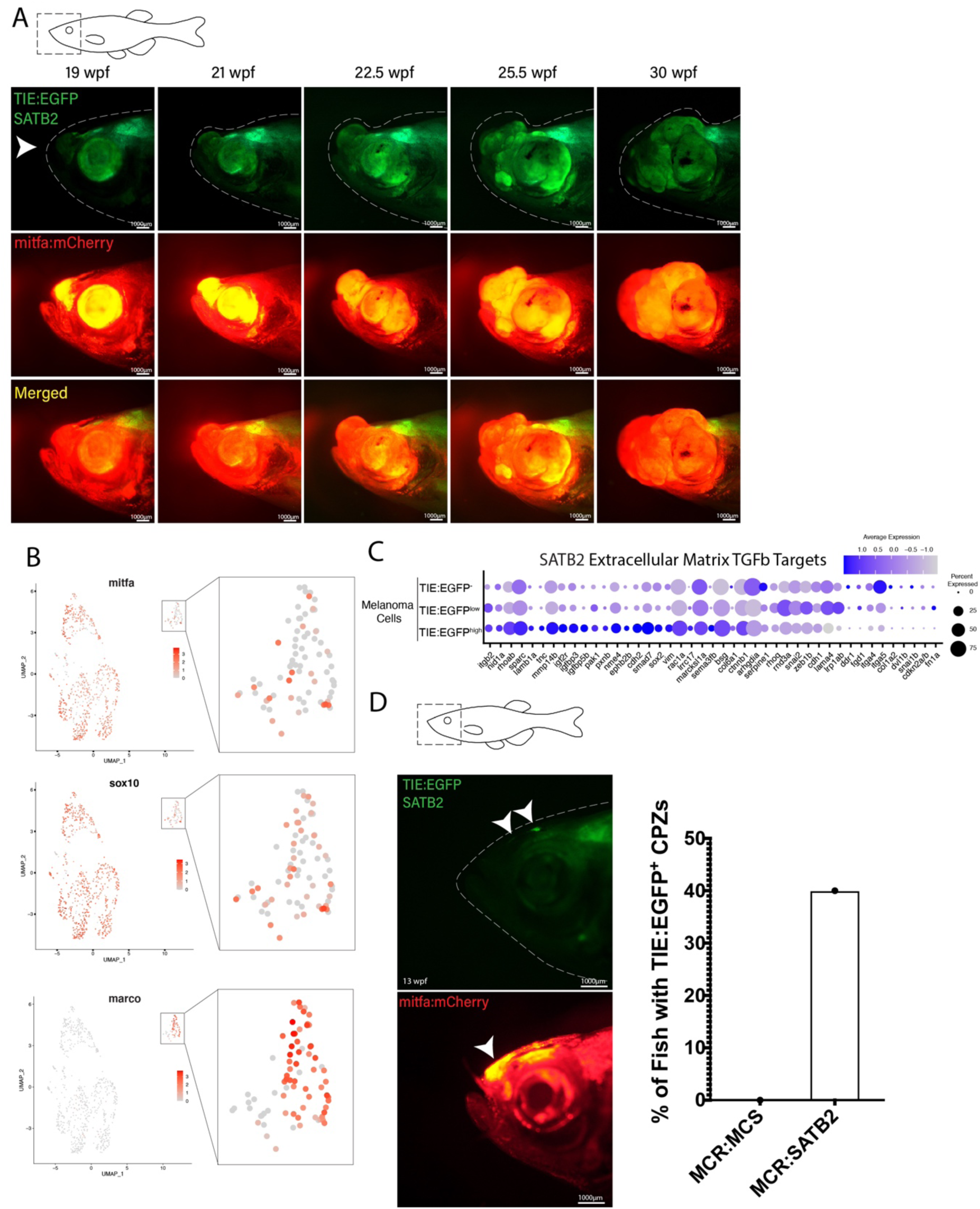
SATB2 expressing melanomas exhibit *TIE:EGFP* expression in early initiating melanomas. (A) Development of a representative tumor overexpressing *MCR:SATB2* in *TIE:EGFP Tg(mitf:BRAF^V600E^); p53^−/−^; mitfa^−/−^* zebrafish. Arrowhead indicates *TIE:EGFP^+^* early melanoma before tumor formation. (B) UMAP depicting *mitfa, sox10,* and *marco* expression in clusters identified by SORT-seq of SATB2 expressing tumor in Figure 4A. Inset shows expression of these genes in the macrophage cluster. (C) Dotplot depicting extracellular matrix TGFb target gene expression in *TIE:EGFP^high^, TIE:EGFP^low^*, and *TIE:EGFP^-^* SATB2 expressing melanoma cells. (D) Early initiating melanoma (arrowhead) overexpressing *MCR:SATB2* in *TIE:EGFP;Tg(mitf:BRAF^V600E^); p53^−/−^; mitfa^−/−^* zebrafish. Representative images chosen. 40% of *MCR:SATB2* early melanomas express *TIE:EGFP*, compared to 0% of *MCR:MCS* tumors (quantification on right).

To determine if there is a change in state of the macrophages that have phagocytosed *TIE:EGFP^+^* cells, we subset macrophages from the *SATB2* expressing melanoma and separated macrophages that were *TIE:EGFP^+^* or *TIE:EGFP^-^*based on flow cytometry intensity. TGFb is known to convert macrophages from a pro-inflammatory M1-like to an anti-inflammatory M2-like state which can be characterized by gene expression. In the zebrafish these include M1 markers *acod1, tnfa, csf3a/b,* and *socs3b*, as well as M2 markers *mrc1b, vegfaa, alox5ap,* and *marco*^44–51^. We found that although *TIE:EGFP^+^* macrophages are not clearly polarized, they express a combination of M1-like and M2-like marker genes (Figure S5B).

To understand the transcriptional differences between *TIE:EGFP^+^* and *TIE:EGFP^-^* cells, *SATB2* expressing melanoma cells were divided into *TIE:EGFP^-^, TIE:EGFP^low^,* and *TIE:EGFP^high^* cells based on flow cytometry intensity. These *MCR:SATB2* results confirmed what was observed in the *TIE:EGFP^+^ MCR:MCS* tumor. We found that chronic extracellular matrix TGFb target genes were up-regulated and the top down-regulated pathways by GSEA again were interferon alpha and gamma (Figure 4C and S5C). Down-regulation of interferon suggests that the TGFb positive melanoma cells are likely evading the immune system. Finally, 40% of the *MCR:SATB2* fish with melanomas had *TIE:EGFP^+^* expressed in early melanoma lesions. These regions are *mitfa:mCherry* high, indicating a proliferation of melanoma cells, but do not extend off the body plane. This is compared to 0% of fish injected with *MCR:MCS* empty vector having *TIE:EGFP^+^*early melanomas (Figure 4D). This shows that overexpression of SATB2 leads to acceleration of TGFb response in melanoma and suggests patients with this amplification may be more susceptible to TGFb inhibitors.

## Discussion

Here we identified an inducible and specific TGFb zebrafish enhancer reporter and visualized TGFb response over melanomagenesis. *TIE:EGFP* is off in early melanoma but is expressed in most advanced melanomas, and remains on, reading out chronic TGFb signaling. These TGFb reporter responsive cells down-regulate interferon, up-regulate a series of novel chronic TGFb target genes and are preferentially phagocytosed by macrophages. This work identifies a TGFb induced immune response and novel biomarkers of chronic TGFb signaling in melanoma.

*TIE:EGFP* expression often occurs in patches throughout the tumor. This zonal TGFb response could be representative of different clonal populations of cells, which bypassed the tumor suppressive effects of TGFb, or because of gradients in the TGFb morphogen. We have shown that our reporter, generated from human ChIP-seq data, is able to read out chronic TGFb signaling. Most literature on developmental signaling focuses on acute signaling with stimulus for several hours *in vitro*. There are multiple explanations for why *TIE:EGFP* is a chronic TGFb reporter and once turned on, remains activated. Once the *TIE:EGFP* reporter is open and active, it may remain active because chromatin is held in an open state by epigenetic modifications or AP-1 factors. We showed that AP-1 factors are necessary for TGFb induction of our TIE reporter and luciferase assays indicated that loss of AP-1 motifs eliminates basal levels of reporter transcription. AP-1 factors may be holding *TIE:EGFP* chromatin open and therefore potentiating the TGFb response. This may allow for a rapid signaling response upon TGFb activation and suggests that AP-1 inhibitors could disrupt TGFb induction.

The *TIE:EGFP* scRNA-seq data indicates that melanoma cells responding to TGFb for several weeks see down-regulation of acute TGFb target genes. They exhibit up-regulation of 29 chronic TGFb target genes that are more representative of the downstream phenotypes imposed by TGFb signal, such as extracellular matrix genes. Based on the literature these targets degrade extracellular matrix promoting migration^40^. Some of the chronic TGFb targets were identified in one of the few reports of long-term TGFb signaling, in human mammary epithelial cells for 12 or 24 days (Figure 2C and 4C). Prolonged TGFb treatment was found to stabilize EMT, stem cell state, and drug resistance in breast cancer cells^39^. According to TCGA data in cBioPortal a subset of melanoma patients exhibit up-regulation of these genes^52, 53^. In the future this signature could be used as biomarkers to identify patients with chronic TGFb signaling and these patients may benefit from treatment with TGFb inhibitors.

Overexpression of *SATB2* resulted in *TIE:EGFP* expression in early lesions, which was not observed in controls. Cancer cells are thought to circumvent the tumor suppressive effects of TGFb via mutations or epigenetic modifications^17, 27, 54–59^. It is possible that melanomas up-regulating the epigenetic regulator SATB2 may overcome the tumor-suppressive effects of TGFb signaling earlier in tumor development, leading to more aggressive and invasive melanomas^43^. Investigating the activation of TGFb signaling in the context of different patient mutations may provide insight into who may benefit from TGFb inhibitors, allowing for the identification of biomarkers of TGFb inhibitor response. It is expected that aggressive melanoma subtypes like *SATB2*, which activate TGFb very early in melanoma development, would be most responsive to early intervention with TGFb inhibitors.

Our work identified macrophage subclusters that contain low levels of *mitfa* and/or *sox10* expression suggestive of phagocytosis of melanoma cells. The vast majority of these macrophages were identified as *TIE:EGFP^+^* by flow cytometry. Using confocal imaging and flow cytometry analysis we confirmed that macrophages were *TIE:EGFP^+^* because they had preferentially phagocytosed a *TIE:EGFP^+^*positive cell. Interestingly, in the tumor pictured in Figure 3C, there is co-localization of *mpeg:mCherry* and *TIE:EGFP* signal indicating that macrophages may cluster in TGFb positive regions of the tumor. As mentioned above, TGFb can act as a chemoattractant for macrophages and induce an anti-inflammatory M2-like state. Single cell RNA-seq data from the *MCR:SATB2* melanoma, suggests macrophages that phagocytosed a TGFb responding cell are not clearly polarized, expressing transcriptional markers of both M1 and M2 states. This may indicate that a transition is occurring from the M1-like to the M2-like phenotype. In our model, macrophages are attracted to TGFb expressing regions of the melanoma and while phagocytosing in the vicinity of TGFb cytokines begin to transition to an M2 state. M2 macrophages are known to be anti-inflammatory, immunosuppressive, and pro-angiogenic. Alternatively, preliminary evidence comparing the transcriptional signatures of macrophages in the *SATB2* expressing tumor, indicates macrophages phagocytosing *TIE:EGFP^+^* cells have higher expression of cholesterol and fatty acid genes (ie. *abca1a, npc2*) as well as apoptotic genes (ie. *casp3b, caspa*) compared to those that phagocytosed *TIE:EGFP^-^* cells. This may indicate a stress phenotype is being induced in macrophages that phagocytose *TIE:EGFP^+^* cells resulting in death. In this case, death of macrophages over time would allow the tumor cells to evade phagocytosis. This, in conjunction with down-regulation of interferon by *TIE:EGFP^+^*melanoma cells, may lead to immune inactivation within the melanoma. Interferon signaling is also down-regulated in *MCR:SATB2* SORT-seq. Moreover, the interferon target gene*, β-2- microglobulin* (*B2M*) is one of the top down-regulated genes in *TIE:EGFP* expressing melanoma cells (Figure S2B). Loss of B2M, part of the MHC Class I molecules, is a known mechanism of immunotherapy resistance^60^. Together, this data suggests that activation of TGFb in melanoma is immunosuppressive, potentially by way of interferon modulation, and supports the need for more work on combination TGFb and immune checkpoint inhibitors.

## Methods

### Plasmids

*Tgfb-Induced-Enhancer:Beta-globin-minimal-promoter-EGFP:pA pDestTol2pA2* was cloned by PCR amplifying the enhancer (chr1:22747452-22747734) in Figure 1A using A375 human melanoma gDNA (Forward primer: TTCTTTGTCATCCTGGTAGAGCAAATCGAG, Reverse primer: GACAGGTCGCACCTGAGTCC)(Advantage® 2 PCR Kit, Clontech #639207, Kyoto, Japan). PCR product was gel purified (Qiagen #28604, Hilden, Germany) and cloned into a pENTR-5’TOPO vector (Invitrogen #45-0711, Waltham, MA, USA). This 5’ entry vector was Gateway recombined upstream of a mouse *Beta-globin-minimal-promoter-EGFP* middle entry vector, a 3’ entry polyA, and the pDestTol2pA2 backbone (ThermoFisher #12538120, Waltham, MA, USA). *Mitfa:mCherry* was Gateway recombined using the *mitfa* zebrafish promoter and a *mCherry* middle entry plasmid (ThermoFisher #12538120). *Ubi:caSMAD2* was cloned by PCR amplifying human *SMAD2* pDONR221 (DNASU HsCD00045549, Tempe, AZ, USA) using the following primers to insert constitutively active mutations (Forward: CACCATGTCGTCCATCTTGCCATTCACGCCGCC; Reverse=CTATtcCATttcTGAGCAACGCACTGAAGGGGATC)(Sigma 11732641001, St. Louis, MO, USA), gel purifying (Qiagen #28604) and inserting into a pENTR™/D-TOPO™ vector (Invitrogen #45-0218). This middle entry vector was Gateway recombined using a 5’ entry zebrafish ubiquitious promoter (Addgene #2732, Watertown, MA, USA), a 3’ entry pA vector, and a pDestTol2pA2 backbone (ThermoFisher #12538120). *Ubi:caSMAD3* was cloned by PCR amplifying human constitutively active *SMAD3* from *Smad3 pCMV-SPORT6* (Harvard Plasmid Repository #HSCD00339271, Boston, MA, USA)(Sigma 11732641001), gel purifying (Qiagen #28604) and inserting into a pENTR™/D-TOPO™ vector (Invitrogen #45-0218). This middle entry vector was Gateway recombined using a 5’ entry zebrafish ubiquitious promoter, a 3’ entry pA vector, and a pDestTol2pA2 backbone (ThermoFisher #12538120). *Ubi:BFP* was cloned via Gateway recombination using a 5’ entry zebrafish ubiquitious promoter, a 3’ entry pA vector, and a pDestTol2pA2 backbone (ThermoFisher #12538120). *Mitfa:BFP* was cloned via Gateway recombination using a 5’ entry zebrafish *mitfa* promoter, a 3’ entry pA vector, and a pDestTol2pA2 backbone (ThermoFisher #12538120). *MCR:SATB2* and *MCR:MCS* are from Fazio et al ^43^. *MCR:BRAF^V600E^* (Addgene #118846) and *2xU6:p53/tyr gRNA mitfa:Cas9* (Addgene #118844) are published (*tyr* gRNA= GGACTGGAGGACTTCTGGGG; *p53* gRNA= GGTGGGAGAGTGGATGGCTG)^61^.

### Reporter line creation

TIE:EGFP reporter fish were created by injecting TIE:Beta-globin-minimal-promoter-EGFP:pA pDestTol2pA2 at 6ng/µL into single cell Tg(mitf:BRAF^V600E^); p53^−/−^; mitfa^−/−^ zebrafish embryos (referred to as TIE:EGFP;Tg(mitf:BRAF^V600E^); p53^−/−^; mitfa^−/−^)^62^. F0s were screened at 5dpf for EGFP. At approximately 2 months post fertilization, F0s were crossed to Tg(mitf:BRAF^V600E^); p53^−/−^; mitfa^−/−^ fish and F1s were screened as embryos for EGFP. EGFP^+^ fish were raised to adulthood.

### Electroporation

Electroporation protocols were adapted from Callahan et al.^63^. *Tg(mitf:BRAF^V600E^); p53^−/−^; mitfa^−/−^* zebrafish were anesthetized using 4% MS-222 (Pentair, TRS1, Minneapolis, MN, USA). Zebrafish were subcutaneously injected with 2µL mix using a Hamilton syringe (Hamilton #80300, Reno, NV, USA) anterior to the dorsal fin. The following amounts of vectors were injected into each fish: 200ng *TIE:Beta-globin-minimal-promoter-EGFP:pA pDestTol2pA2,* 400ng *ubi:caSMAD2:pA*, 400ng *ubi:caSMAD3:pA*, 666ng *ubi:BFP:pA*, and 266ng *Tol2 pCS2FA- transposase*^64^. DNA was prepped using Qiagen HiSpeed Plasmid Maxi Kit (Qiagen #12662). Zebrafish were then electroporated with a BTX ECM 830 machine (BTX #45-0662 Holliston, MA, USA) using the following parameters: LV Mode; 45V; 60ms pulse length; 5 pulses, 1 second interval. The cathode paddle was placed on the side of the fish that was injected. Electroporated fish were imaged approximately one week post electroporation using a Nikon SMZ18 Stereomicroscope (Nikon, Tokyo, Japan). To quantify TIE activity, we imaged each electroporated fish, quantified GFP intensity, and divided by the area of BFP to account for variation in electroporation efficiency.

### Imaging

Zebrafish were anesthetized using 4% MS-222 (Pentair, TRS1) and imaged using a Nikon SMZ18 Stereomicroscope or a Nikon C2si Laser Scanning confocal. Z-stacks were aligned and images minimally processed using Imaris (RRID:SCR_007370) or ImageJ software (RRID:SCR_003070). Confocal images were taking using a 10x objective.

### Inhibitor treatment

*TIE:EGFP*;*Tg(mitf:BRAF^V600E^); p53^−/−^; mitfa^−/−^* F1s were crossed and F2 embryos were screened for EGFP, dechorionated at 24 hpf, and placed in a 24 well plate. Embryos were treated with either E3 zebrafish water, DMSO vehicle control (Sigma #D2650), 50µM or 100µM SB 431542 hydrate (Sigma # S4317-5MG). 24 hours post treatment embryos were imaged using a Nikon SMZ18 Stereomicroscope (1x objective; 3x zoom; 7ms white light and 300ms GFP exposure used for all conditions).

### Melanoma generation

Melanocyte development is conserved between zebrafish and mammals^65^. The master regulator of the melanocyte lineage, MITF, is conserved in zebrafish (*mitfa*) and required for melanocyte development^66^. Expression of *mitfa:BRAFV600E* together with a homozygous p53 missense mutation, leads to development of zebrafish melanomas^62^. A *mitfa*^−/−^ mutation in *Tg(mitfa:BRAFV600E);p53^−/−^* fish prevents melanocyte development and spontaneous melanoma formation. Melanocyte development can then be rescued via injection of MiniCoopR (MCR), a transposon-based vector that contains a *mitfa* minigene alongside a candidate oncogene driven by the *mitfa* promoter^67^. *TIE:EGFP; Tg(mitf:BRAF^V600E^); p53^−/−^; mitfa^−/−^; MCR:MCS; mitfa:mCherry; tyr^-/-^* zebrafish embryos were generated by injecting single cell *TIE:EGFP;Tg(mitf:BRAF^V600E^); p53^−/−^; mitfa^−/−^* zebrafish embryos with *MiniCoopR Multiple Cloning Site (MCR:MCS)* at 20ng/µL, *mitfa:mCherry* at 10ng/µL, tyr gRNA (GGACTGGAGGACTTCTGGGG) at 10ng/µL, Cas9 protein (PNA Bio CP02) at 50ng/µL, and Tol2 mRNA at 20ng/µL^67^. Injection of the MCR vector, which contains a *mitfa* minigene mosaically rescues melanocytes in *Tg(mitf:BRAF^V600E^); p53^−/−^; mitfa^−/−^* zebrafish. In the absence of a candidate oncogene the *mitfa* promoter is followed by an empty multiple cloning site (*MCR:MCS*). Tyrosinase gRNA was created by annealing a *tyr* oligo template (CCTCCATACGATTTAGGTGACACT ATAGGACTGGAGGACTTCTGGGGGTTTTAGAGCTAGAAATAGCAAG) to a constant oligonucleotide (AAAAGCACCGACTCGGTGCCACTTTTTCAAGTTGATAACGGACTAGCCTTATTTTAA CTTGCTATTTCTAGCTCTAAAAC). The annealed oligo was filled in using T4 DNA polymerase (New England BioLabs, M0203S, Ipswich, MA, USA), PCR amplified, gel purified, transcribed (MEGAscript T7/SP6 ThermoFisher Scientific, AM1333), and cleaned up (Zymo Research, R2051, Irvine, CA, USA).

### Zebrafish work

All zebrafish (*Danio rerio*) work was performed in accordance with the Guide for the Care and Use of Laboratory Animals of the National Institutes of Health. Animal research protocols were approved by the Institutional Animal Care and Use Committee of Boston Children’s Hospital, Protocol #20-10-4254R. All zebrafish work operated according to the guidelines of the Institutional Animal Care and Use Committee of Boston Children’s Hospital.

### SORT-seq

Tumors were excised and dissociated for 30 minutes with occasional chopping using 0.075mg/mL liberase (Sigma #5401119001) in DMEM (Gibco #11965-092, Waltham, MA, USA) with 1% Penstrep (Corning #30-002-CI, Corning, NY, USA). Casper zebrafish skin was used to exclude autofluorescence when setting gates^68^. A *mitfa:mCherry^+^/EGFP^-^* tumor from a TIE:EGFP *Tg(mitf:BRAF^V600E^); p53^−/−^; mitfa^−/−^* fish or *mitfa:mCherry;crestin:EGFP;tyr^-/-^* zebrafish skin was used to set the gates for mCherry intensity. *Ubi:GFP* zebrafish skin was used to set the gate for GFP intensity. Dissociated samples were filtered through a 40µm filter and resuspended in FACs buffer (PBS/ 10%FBS/1% Penstrep) before filtering through a FACs tube (Corning #352235). Single cells were sorted into 384-well-cell-capture plates containing barcoded primers from Single Cell Discoveries (https://www.scdiscoveries.com/) using a BD FACS ARIA II sorter. SYTOX™ was used as a live/dead marker (ThermoFisher #S34857). Library preparation and Illumina sequencing was performed by Single Cell Discoveries^38^. Sort-seq data are demultiplexed and aligned to zebrafish Ensembl GRCz11 annotation using scruff R packages with the following parameter (bcStart = 7, bcStop = 14, bcEdit = 1, umiStart = 1, umiStop = 6, keep = 60)^69^. The 384 sort-seq barcodes are downloaded from https://github.com/anna-alemany/transcriptomics/blob/master/mapandgo/bwamap/bc_celseq2.tsv.

Analysis was completed in R Studio using Seurat (RRID:SCR_016341)(min.features= 200; 600<nFeatureRNA<10000; percent.mt<10; obj.resolution= 0.2; GFPhigh>4000)^70–73^. Batch correction was performed using FindVariableFeatures and FindIntegrationAnchors, nfeatures=15000. Pathway analysis was conducted using Gene Set Enrichment Analysis (GSEA; RRID:SCR_003199) version 4.1.0^74, 75^. Zebrafish genes were converted to human using DIOPT Ortholog Finder version 8.5 (RRID:SCR_021963) and the best match was used for GSEA analysis^76^.

### Confocal Imaging

Mpeg1:mCherry Casper zebrafish (roy^-/-^; mitfa^-/-^) were crossed to TIE:EGFP;Tg(mitf:BRAF^V600E^); p53^−/−^; mitfa^−/−^ fish and embryos were injected with 2x U6:p53/Tyr gRNA mitfa:Cas9, mitfa:BRAF^V600E^ MCR, and mitfa:BFP at 10ng/µL, along with Tol2 mRNA at 20ng/µL^61^. Embryos were grown to adulthood and TIE:EGFP^+^ tumors were imaged using Nikon C2si Laser Scanning confocal at 10X. Z-stacks were aligned and images minimally processed using Imaris or ImageJ software.

### Flow Analysis

Tumors were excised and dissociated for 30 minutes with occasional chopping using 0.075mg/mL liberase (Sigma #5401119001) in DMEM (Gibco #11965-092) with 1% Penstrep (Corning #30-002-CI). Casper zebrafish skin was used to exclude autofluorescent cells^68^. *Mpeg:mCherry^+^* and *Ubi:GFP^+^* zebrafish skin was used to set the gates for mCherry and GFP intensity, respectively. A *mitfa:BFP^+^*patch of skin as well as *Flk:BFP^+^* skin was used to set the gates for BFP intensity. Dissociated samples were filtered through a 40µm filter and resuspended in FACs buffer (PBS/10%FBS/1% Penstrep) before filtering through a FACs tube (Corning #352235). DRQ7™ was used as a live/dead marker (Abcam #ab109202, Cambridge, UK). Cells were analyzed using a BD FACS Aria II 5 Laser System. Two technical replicates of 1 million cells were sorted from each tumor, for a total of 2 million cells per tumor. Data was processed using FlowJo version 10.8.1 (RRID:SCR_008520). Debris, doublets, and autofluorescent cells were removed from the analysis and viable cells were separated into *TIE:EGFP^-^* and *TIE:EGFP^+^*.

### Human and Zebrafish Melanoma Cell Culture and Treatment

Human melanoma A375 and zebrafish melanoma ZMEL1 cells were grown in filter sterilized DMEM (Gibco #11965-092) supplemented with 10% heat-inactivated FBS (Gemini # 900-108), 1% PenStrep (Corning #30-002-CI), and 1% Glutamine (ThermoFisher # 25030164). A375 cells were obtained from ATCC (RRID:CV-CL_0132) and grown at 37°C, 5% CO_2_. Cell identity was confirmed by fingerprint every 2 years and tested for mycoplasma approximately every 2-4 weeks using Lonza’s second generation myco PLUS kit. ZMEL1 cells were grown at 28°C, 5% CO_2_ and tested for mycoplasma approximately every 2-4 weeks^77^.

Human recombinant TGFB1 (R&D 240-B-002, Minneapolis, MN, USA) was reconstituted at 20µg/mL in sterile 4mM HCl containing 1mg/mL BSA according to manufacturer’s instructions. To activate the TGFb pathway, cells were serum starved for 2 hours, then treated with 10ng/mL human recombinant TGFB1 or 4mM HCl containing 1 mg/mL BSA vehicle control for an additional two hours.

### RNA-sequencing

RNA-seq was performed in triplicate using A375 or ZMEL1 melanoma cells. RNA was collected from adherent cells using the Qiagen RNeasy Plus Mini Kit (Qiagen #74134). RNA quality was confirmed using a Fragment Analyzer. One microgram of RNA was ribodepleted using NEBNext rRNA Depletion Kit (NEB #E6310). Ribodepleted RNA was fragmented, reverse transcribed, and library prepped (NEB #E7530, NEB #E7335). Samples were sequenced on an Illumina Hi-Seq 4000 sequencer. Quality control of RNA-Seq datasets was performed by FastQC (https://www.bioinformatics.babraham.ac.uk/projects/fastqc/)(RRID:SCR_014583) and Cutadapt (RRID:SCR_011841) to remove adaptor sequences and low quality regions ^78^. The high-quality reads were aligned to Ensembl build version GRCh38 of human genome or zebrafish Ensembl GRCz11 annotation (RRID:SCR_002334) using Tophat 2.0.11 (RRID:SCR_013035) without novel splicing form calls^79^. Transcript abundance and differential expression were calculated with Cufflinks 2.2.1 (RRID:SCR_014597)^80^. FPKM values were used to normalize and quantify each transcript; the resulting list of differentially expressed genes are filtered by log fold change and q-value. Pathway analysis was conducted using Gene Set Enrichment Analysis (GSEA) version 4.1.0 Hallmark gene sets^74, 75^. DEseq2 (RRID:SCR_015687) was used to create differential expression heatmaps and volcano plots. Read counts less than 10 were excluded.

### ChIP-sequencing

A375 cells were fixed directly in 15cm plates with 11% formaldehyde and collected by scraping. Approximately 100,000,000 cells were used per condition. Cells were lysed using lysis buffers with protease inhibitors (Roche #05056489001, Basel, Switzerland) and sonicated such that fragmented chromatin was 200-300bp long. Optimal chromatin length was confirmed by gel electrophoresis. Prior to antibody addition, 50µL chromatin was collected for input sample. The remaining sonicated chromatin was incubated overnight at 4°C with 10µg antibody attached to Dynabeads (Invitrogen #10004D). Samples (including inputs) were washed with wash buffers and eluted for 6 hours at 70°C, treated with RNaseA (Sigma #R4642) and ProteinaseK (Life Technologies #AM2546, Carlsbad, CA, USA), and purified using Zymo ChIP DNA Concentrator kit (Genesee Scientific #11-379C, San Diego, CA, USA). Libraries were end repaired (VWR #ER81050, Radnor, PA, USA), polyA tailed (Invitrogen #18252-015, NEB #M0212L), adaptor ligated (NEB #E7335), size selected using Ampure XP beads (Beckman Coulter #A63881, Brea, CA, USA, Life Technologies #12027), and PCR amplified (NEB #M0531). Libraries were run on an Illumina Hi-Seq 4000 sequencer. The following antibodies were used: H3K27ac (Abcam #4729; RRID:AB_2118291), SMAD2/3 (Abcam #202445), MITF (Sigma #HPA003259; RRID:AB_1079381), ATF3 (Abcam #207434; RRID:AB_2734728), JUNB (CST #3753, Danvers, MA, USA; RRID:AB_2130002). Using HOMER analysis (RRID:SCR_010881) we confirmed that JUNB and ATF3 binding motifs were present under their respective ChIP peaks (ATF3, p=1e-4983) (JUNB, p=1e-15697). All ChIP-Seq datasets were aligned to Ensembl build version GRCh38 of the human genome using Bowtie2 (version 2.2.1; RRID:SCR_016368) with the following parameters: --end-to-end, -N0, -L20 ^81^. MACS2 version 2.1.0 (RRID:SCR_013291) peak finding algorithm was used to identify regions of ChIP-Seq peaks, with a q-value threshold of enrichment of 0.05 for all datasets^82^. Uropa (Universal RObust Peak Annotator) is utilized to annotate ChIP-seq peaks to neighboring genes according to Ensembl gene annotation^83^. The parameters are defined as proximal promoter: 500 bp upstream - 50 bp downstream of TSS; distal promoter: 2k bp upstream - 500 bp downstream of TSS; enhancer: 100k bp from TSS. The genome-wide transcription factor SMAD2/3, JUNB, ATF3 occupancy profile figures were generated by deeptools2 according to two computation modes^84^. In the reference-point mode, a set of genomic positions (e.g. the center of ChIP peak) are used as anchor point, 2 kb upstream and downstream of these position are plotted in the profile figure. HOMER analysis was performed to confirm transcription factor binding under peaks ^85^. The hg19 genome was used with a random set of background peaks for motif enrichment.

### Luciferase Assays

Firefly luciferase reporter constructs (pGL4.24) were created by cloning the full and mutated TGFb enhancers upstream of the minimal promoter using BglII and XhoI sites (see Appendix Figure 1 for sequences). A375 cells were plated in opaque-walled 96-well plates (ThermoFisher #136101) and approximately 5,000 cells were co-transfected with 100ng firefly and 10ng Renilla luciferase plasmids using Lipofectamine 3000 (Invitrogen #L3000008). After 48 hours cells were serum starved for 2 hours and treated with 10ng/mL TGFB1 or 4mM HCl containing 1mg/mL BSA vehicle control for an additional 2 hours. Firefly and Renilla luciferase were then measured using the Dual-Glo Luciferase Assay (Promega #E2920, Madison, WI, USA) according to the manufacturer’s instructions. Luminescence was read on a Synergy Neo plate reader and the ratio of firefly to Renilla luminescence was calculated. Empty firefly luciferase vector was used as a negative control and Renilla luciferase was used as control for transfection efficiency. Experiments were performed in biological triplicate with three technical replicates each.

### Statistics

To calculate significance of electroporation and flow analysis assays, a 2-tailed unpaired t test with Welch’s correction was performed using GraphPad Prism version 9.0.2 (RRID:SCR_002798) for Mac, GraphPad Software, San Diego, California USA, www.graphpad.com. To calculate significance of luciferase assays, a 2-way multiple comparison ANOVA test was performed using GraphPad Prism version 9.0.2 for Mac, GraphPad Software, San Diego, California USA, www.graphpad.com.

### Material availability

The *Tgfb-Induced-Enhancer:Beta-globin-minimal-promoter-EGFP:pA pDestTol2pA2* plasmid will be made available on Addgene and the *TIE:EGFP* zebrafish reporter line is available upon request. Sequencing datasets can be accessed on GEO # GSE213360. Reviewers can access data here: https://www.ncbi.nlm.nih.gov/geo/query/acc.cgi?acc=GSE213360; token: ifcbckwqddgzjgx.

## Acknowledgements

We would like to thank Christian Lawrence, Kara Maloney, Shane Hurley, Lauren McKay, Li- Kun Zhang, and Andrew Kowalczyk for fish care throughout this study. We would also like to thank the Boston Children’s Hospital Flow Cytometry Research Facility for assistance with FACS experiments, as well as Single Cell Discoveries for processing SORT-seq samples. The authors would like to thank their colleagues for discussion and reading of this manuscript. Figures were created with Biorender.com. L.I.Z. is a Howard Hughes Medical Institute Investigator. Additional funding for this work was provided by the Human Frontier Science Program (HFSP LT000494/2020-L, to C.S.B.).

## Competing interests

L.I.Z. is a founder and stockholder of Fate Therapeutics, CAMP4 Therapeutics, Amagma Therapeutics, Scholar Rock, and Branch Biosciences. He is a consultant for Celularity and Cellarity.

---

**Supplemental Figure 1.**
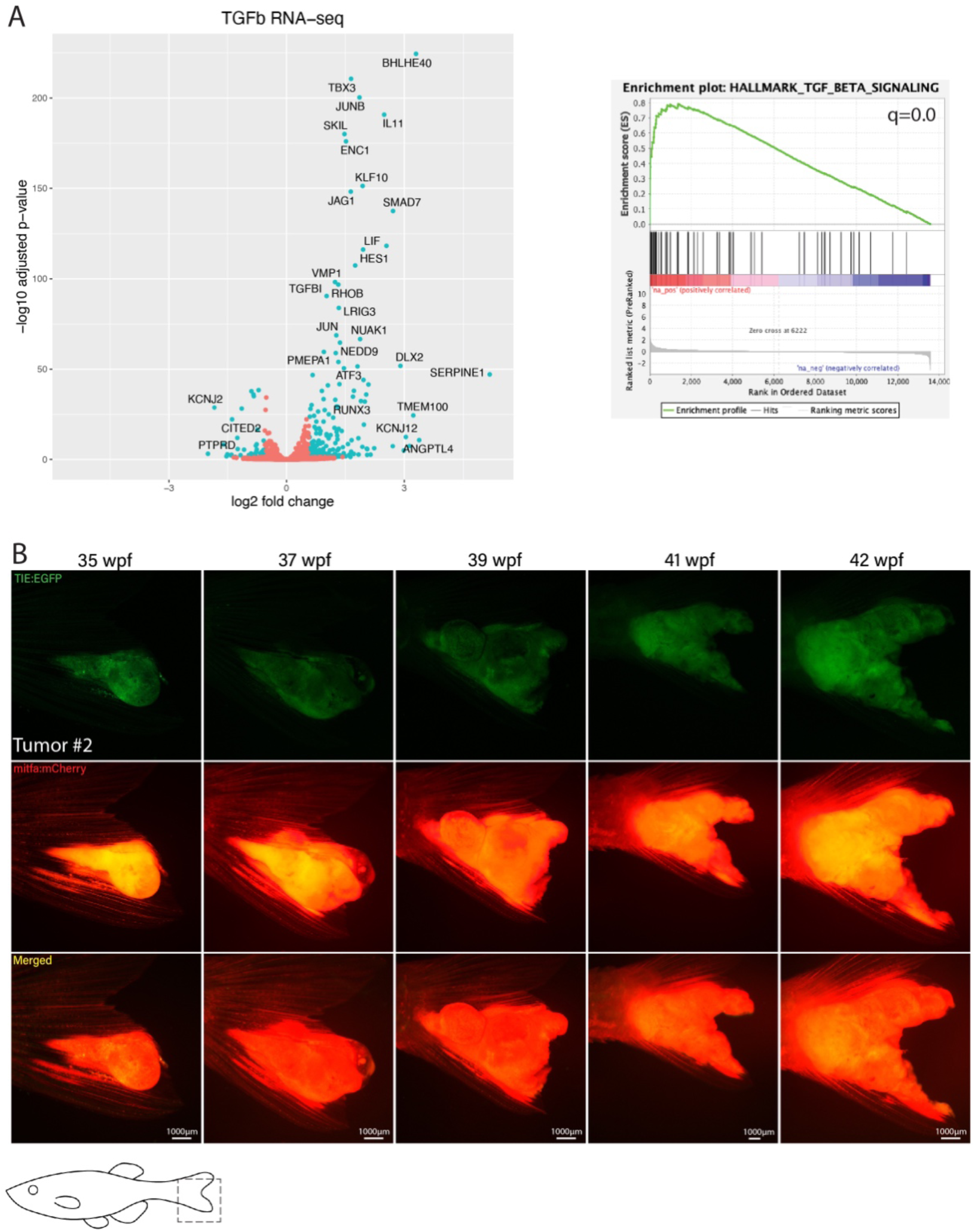
Treatment with human recombinant TGFB1 activates the TGFb pathway in human melanoma cells. (A) Left: RNA-sequencing indicated that following a 2-hour treatment with 10ng/mL TGFB1, 223 genes were significantly up-regulated (q<0.05) and 94 genes were down-regulated. This includes typical TGFb target genes such as SMAD7, JUNB, and PMEPA1. Volcano plot indicating top up and down-regulated genes upon TGFB1 treatment. Right: Hallmark gene set enrichment analysis (GSEA) of genes ranked by log2 fold-change (log2fc) confirmed that TGFb was the top up-regulated pathway (q=0). RNA-seq was performed in triplicate. (B) *TIE:EGFP^+^* melanoma #2 expressing *MCR:MCS*. *TIE:EGFP* expression across melanomagenesis. Representative images shown.

**Supplemental Figure 2.**
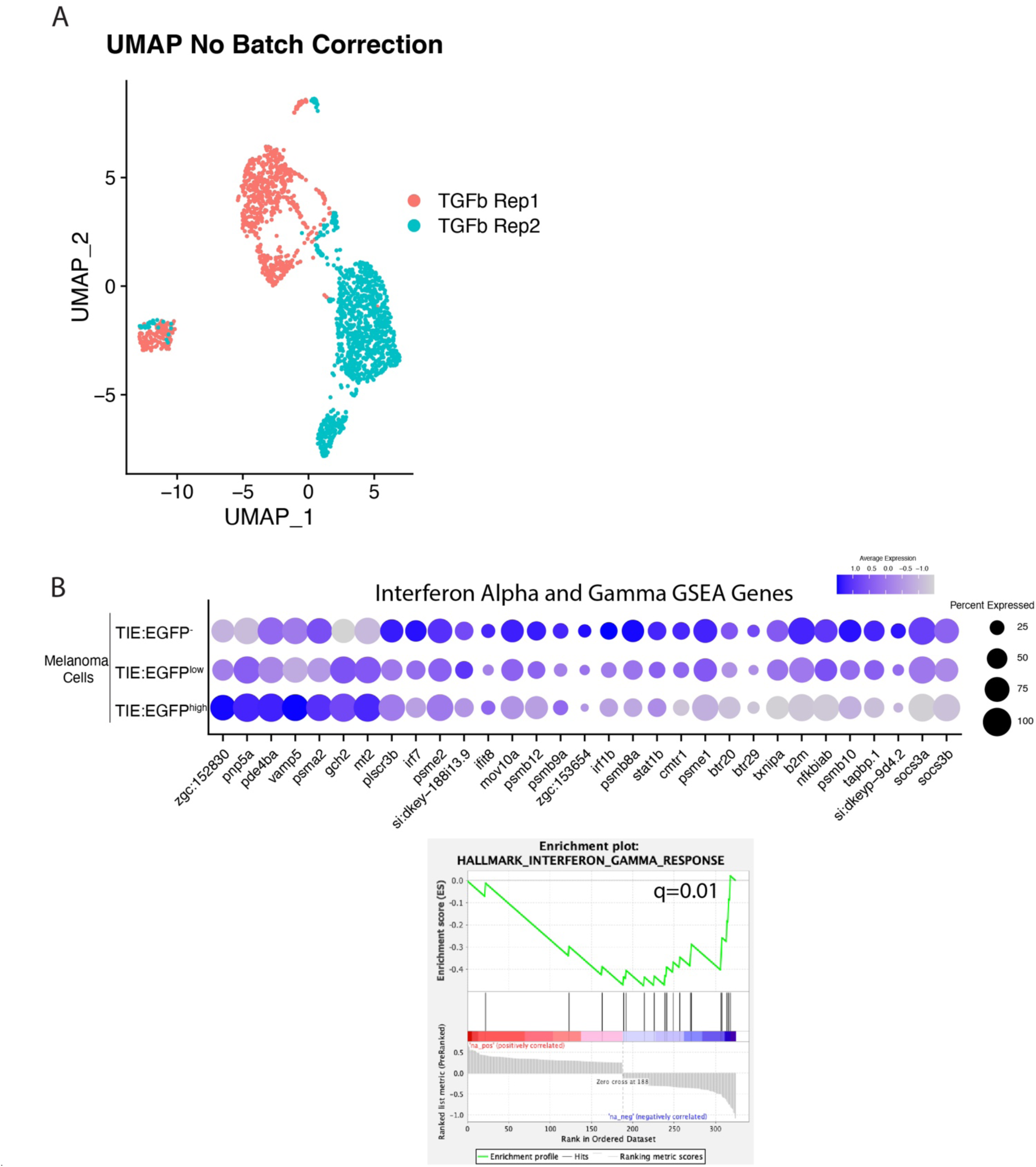
(A) UMAP with no batch correction of 2 *MCR:MCS* replicates combined in Figure 2A. Approximately 2,256 *TIE:EGFP^+^*cells and 752 *mitfa:mCherry^+^* cells were sorted for scRNA-seq. (B) Dotplot depicting interferon target gene expression in *TIE:EGFP^high^, TIE:EGFP^low^*, and *TIE:EGFP^-^* melanoma cells. GSEA enrichment plots reveal down-regulated interferon gamma (q=0.01) in *TIE:EGFP^+^*melanoma cells (bottom).

**Supplemental Figure 3.**
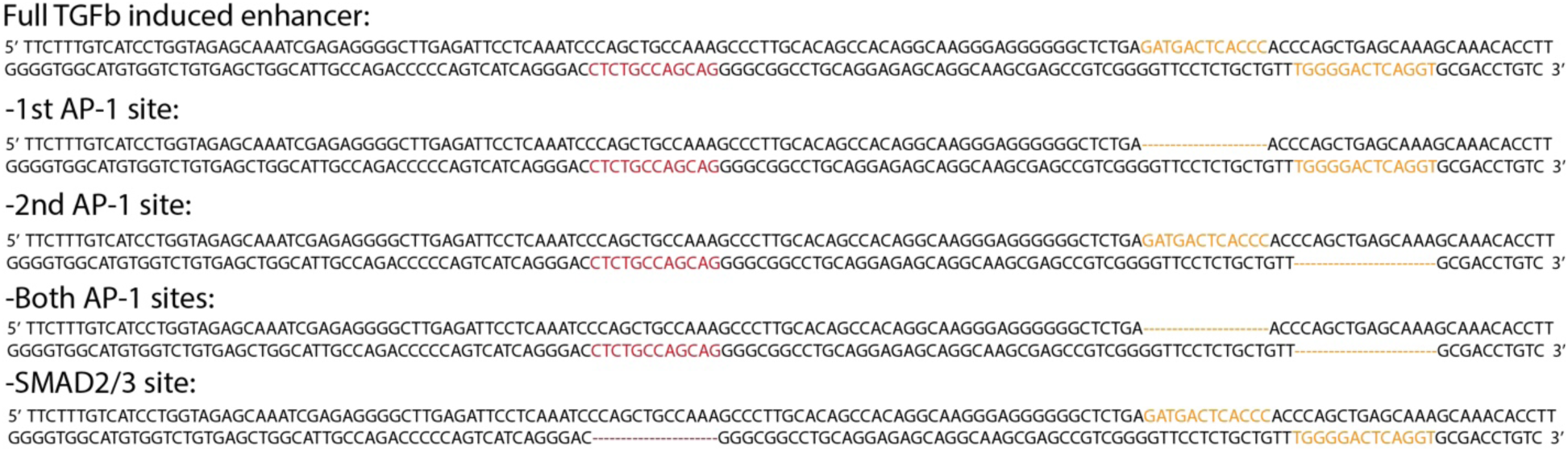
Mutated TGFb inducible enhancer sequences used for luciferase assays in Figure 4C. AP-1 motifs are in orange, SMAD2/3 motif is in red.

**Supplement Figure 4.**
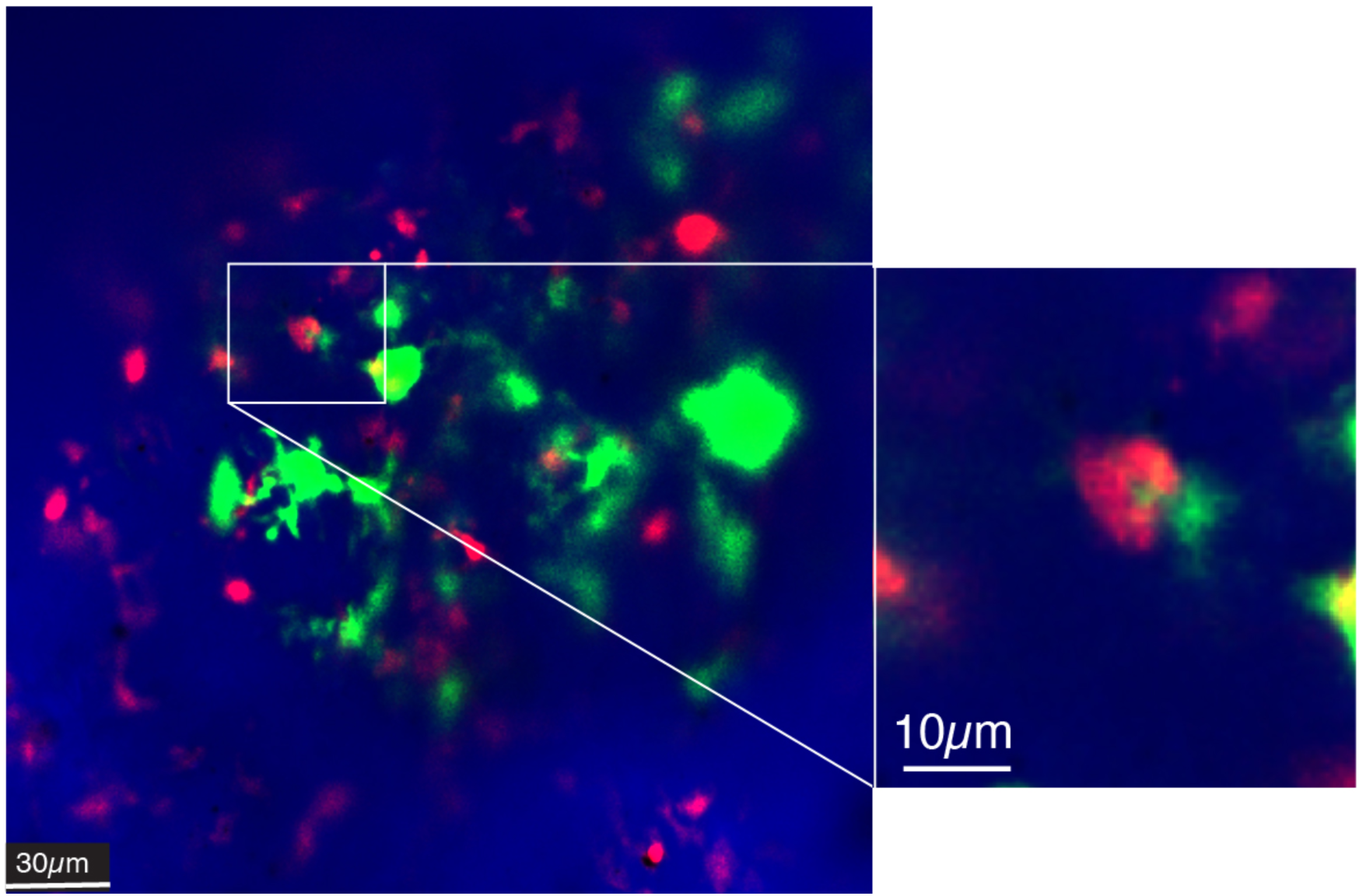
Example of a macrophage actively phagocytosing a *TIE:EGFP^+^* cell in a melanoma.

**Supplement Figure 5.**
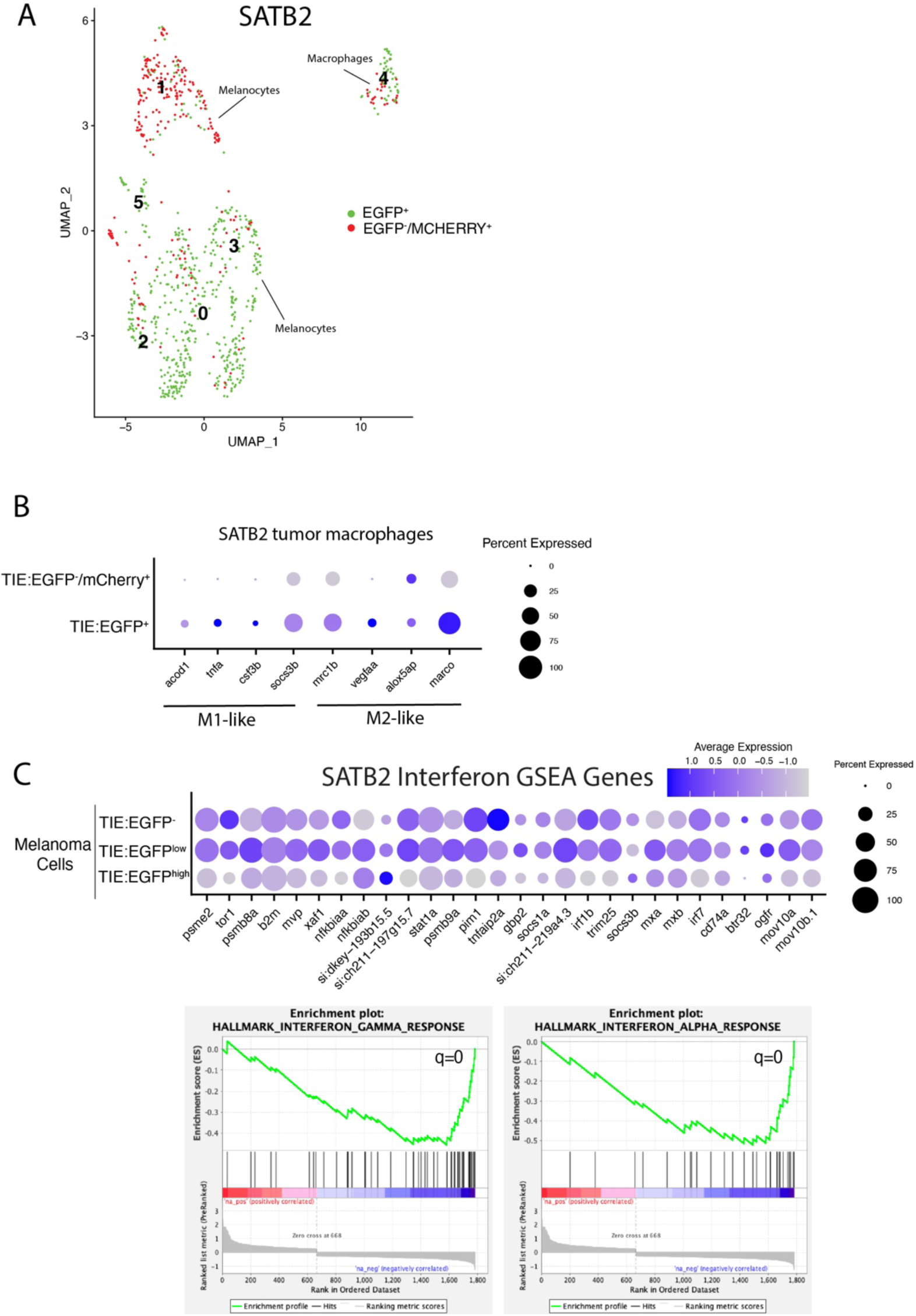
Single cell RNA-seq identifies *MCR*:*SATB2 TIE:EGFP*^+^ macrophages and melanoma cells. SATB2 TGFb expressing melanoma cells down-regulate interferon. (A) UMAP depicting 6 cell clusters identified by SORT-seq (left). *TIE:EGFP^+^* and *TIE:EGFP^-^* cells are identified based on flow cytometry. Melanoma cells were identified as being *mitfa* and *sox10* positive, while macrophages are *marco* positive. (B) Dotplot depicting M1-like and M2-like transcriptional marker expression in *TIE:EGFP^+^*and *TIE:EGFP^-^* macrophages from SATB2 TGFb expressing melanoma. (C) Dotplot depicting expression of interferon alpha and gamma GSEA pathway genes in *TIE:EGFP^high^, TIE:EGFP^low^,* and *TIE:EGFP^-^* melanoma cells. GSEA enrichment plots reveal down-regulated interferon alpha (q=0) and gamma (q=0.0) in *TIE:EGFP^+^* melanoma cells (bottom).

